# Triangulation of microbial fingerprinting in anaerobic digestion reveals consistent fingerprinting profiles

**DOI:** 10.1101/2021.05.28.446109

**Authors:** Jo De Vrieze, Robert Heyer, Ruben Props, Lieven Van Meulebroek, Karen Gille, Lynn Vanhaecke, Dirk Benndorf, Nico Boon

## Abstract

The anaerobic digestion microbiome has been puzzling us since the dawn of molecular methods for mixed microbial community analysis. Monitoring of the anaerobic digestion microbiome can either take place *via* a non-targeted holistic evaluation of the microbial community through fingerprinting or by targeted monitoring of selected taxa. Here, we compared four different microbial community fingerprinting methods, *i.e.*, amplicon sequencing, metaproteomics, metabolomics and cytomics, in their ability to characterise the full-scale anaerobic digestion microbiome. Cytometric fingerprinting through cytomics reflects a, for anaerobic digestion, novel, single cell-based approach of direct microbial community fingerprinting by flow cytometry. Three different digester types, *i.e.*, sludge digesters, digesters treating agro-industrial waste and dry anaerobic digesters, each reflected different operational parameters. The α-diversity analysis yielded inconsistent results, especially for richness, across the different methods. In contrast, β-diversity analysis resulted in comparable profiles, even when translated into phyla or functions, with clear separation of the three digester types. In-depth analysis of each method’s features *i.e.*, operational taxonomic units, metaproteins, metabolites, and cytometric traits, yielded certain similar features, yet, also some clear differences between the different methods, which was related to the complexity of the anaerobic digestion process. In conclusion, cytometric fingerprinting through flow cytometry is a reliable, fast method for holistic monitoring of the anaerobic digestion microbiome, and the complementary identification of key features through other methods could give rise to a direct interpretation of anaerobic digestion process performance.

## 1. Introduction

The anaerobic digestion (AD) microbiome has been one of the most studied engineered ecosystems since the dawn of molecular methods. In the 1990s and early 2000s, taxa identification took place via labour-intensive techniques, such as clone libraries (Godon et al., 1997; Mladenovska et al., 2003), while semi-quantification could be achieved through fluorescent *in situ* hybridization (FISH) (Raskin et al., 1994). Comprehensive microbial community profiling was performed *via* basic fingerprinting techniques, such as denaturing gradient gel electrophoresis (DGGE) (Liu et al., 2002) or terminal restriction fragment length polymorphism (T-RFLP) (Collins et al., 2003). Even though such methods provided relevant results towards the AD microbiome’s unravelling, the emergence of high-throughput DNA sequencing methods about fifteen years ago (Hugerth and Andersson, 2017; Sogin et al., 2006) was a real game-changer that quickly accelerated the endeavours to characterise the AD microbiome. With 16S rRNA gene amplicon sequencing, the core taxa in AD were identified (Calusinska et al., 2018; Rui et al., 2015; Tao et al., 2020), and their link with operational parameters clarified (De Vrieze et al., 2018, 2015; Mei et al., 2017; Sundberg et al., 2013). The “omics” enabled the transition from identification and fingerprinting towards, amongst others, the complementary estimation of the physiological potential (metagenomics), activity (metatranscriptomics or metaproteomics) and metabolic performance (metabolomics) of the AD microbiome (Hassa et al., 2018; Heyer et al., 2015; Vanwonterghem et al., 2014).

Given the target-specific nature of these different high-throughput methods, *i.e.*, amplicon sequencing and the “omics”, and because of the complexity of natural and engineered microbiomes, more and more studies focused on combining them to answer specific questions (Franzosa et al., 2015), though this often remains challenging (Prosser, 2015). Combining 16S rRNA gene amplicon sequencing, metagenomics and metatranscriptomics revealed a high activity of methanogenic archaea compared to their low (relative) abundance in AD (Zakrzewski et al., 2012). These results were confirmed through the combination of metagenomics and metatranscriptomics, where it was highlighted that a high abundance of certain taxa does not necessarily relate to high activity in AD (Maus et al., 2016). In activated sludge, the same combination of “omics” allowed for determining the identity and activity of taxa containing antibiotic resistance genes (Liu et al., 2019). Through the coupling of metagenomics and metaproteomics, the high metabolic activity of methanogenic archaea in AD was confirmed (Hanreich et al., 2013). In another study, a high degree of similarity, both in terms of β-diversity and taxon identification, was observed between 16S rRNA gene amplicon sequencing and metaproteomics (Heyer et al., 2016). A combination of metagenomics and metabolomics made it possible to link microbial community composition and related operational parameters with certain metabolites (Beale et al., 2016). These studies provided novel insights into the microbiome in AD on different levels, both for overall fingerprinting and targeted analyses.

In the last decade, the development of a novel approach that enables single-cell measurements of cytometric traits through flow cytometry (cytomics) allowed the possibility to monitor the microbiome at a new level through a so-called “cytometric fingerprint” (De Roy et al., 2012; Props et al., 2016). In contrast to the labour-intensive and time-consuming nature of other levels of fingerprinting, the cytometric fingerprint in aquatic matrices can be obtained in a matter of minutes to hours, and can allow absolute quantification of cells (Besmer et al., 2017, 2014; Props et al., 2017). The combination and high similarity in cytometric fingerprinting and 16S rRNA gene amplicon sequencing in a natural bacterioplankton community (Props et al., 2018) and for drinking water monitoring (Prest et al., 2014) indicate this method’s potential.

Despite its unique features, this high-throughput optic-based method has received limited attention in engineered microbial processes, mainly related to the complex matrix of, *e.g.*, anaerobic digestate and waste activated sludge (Brown et al., 2019). Thus far, flow cytometry has been used in AD for archaea monitoring through the F_420_ co-factor (Lambrecht et al., 2017), for evaluation of microbial community dynamics and the formation of sub-communities (Günther et al., 2018), or for the direct phenotypic monitoring of the microbial community (Dhoble et al., 2016). Hence, cytometric monitoring through flow cytometry of the AD microbial community remains confined to a limited number of studies, and cytometric fingerprinting analyses in AD lack an accurate comparison with current fingerprinting methods at different levels, *i.e.*, amplicon sequencing and the “omics”.

This study’s key objective was to compare three established methods, *i.e.*, 16S rRNA gene amplicon sequencing, metaproteomics, metabolomics, with cytomics, through the concept of triangulation (Lawlor et al., 2016), regarding their characterisation of the microbiome in fullscale AD plants. These methods were compared at the overall, non-targeted fingerprinting level and through targeted analysis, *i.e.*, identification of taxa, metabolites and functionality. We investigated the features (operational taxonomic units (OTUs), metaproteins, metabolites, and cytometric traits) of the fingerprinting methods to illuminate the methods’ different information gain.

## 2. Materials and methods

### 2.1. Sample and data collection

Digestate samples were collected from 63 full-scale AD plants in Belgium in 1 L air-tight containers. The samples were homogenized upon arrival in the laboratory, and three replicate 1.5 mL subsamples were stored at −80°C until DNA (amplicon sequencing), protein (metaproteomics) and metabolite (metabolomics) extraction. A 10 mL subsample was stored at −20°C for volatile fatty acids (VFA) analysis. A 50 mL sample was stored at 4°C for volatile solids (VS), total solids (TS), total ammonia nitrogen (TAN), and cation (Na^+^ and K^+^) analysis. The pH and conductivity of each sample were measured immediately upon arrival in the laboratory. Information concerning the sludge retention time (SRT) and temperature in the digester was obtained directly from each plant operator.

### 2.2. Microbial community analysis

#### 2.2.1. Amplicon sequencing

The DNA was extracted directly from the at −80°C frozen samples using an optimized method (Vilchez-Vargas et al., 2013). The DNA extracts’ quality was validated through a 1% agarose gel electrophoresis and via PCR with the bacterial primers P338 and P518r (Muyzer et al., 1993), following the protocol of Boon et al. (2002). The DNA extracts were normalised to contain 1 ng μL^−1^ DNA, and sent out to LGC Genomics GmbH (Berlin, Germany) for library preparation and sequencing on an Illumina Miseq platform. The amplicon sequencing and data processing were carried out as described in SI1 (S1). A table with the abundance of OTUs (operational taxonomic units) and their taxonomic assignments in each sample was generated (SI2), and absolute singletons were removed. The raw fastq files that served as a basis for the bacterial community analysis were deposited in the National Center for Biotechnology Information (NCBI) database (Accession number SRP321483).

#### 2.2.2. Metaproteomics

Sample preparation for metaproteomics was carried out as described earlier (Heyer et al., 2019), starting from the at −80°C frozen samples. Purified peptide mixtures were analysed by reversed-phase liquid chromatography coupled to a timsTOF™ Pro tandem mass spectrometer (Bruker Daltonik GmbH, Bremen), using a 120 min gradient. For protein identification, the mass spectrometer data were searched with Mascot™ 2.6.1 (Matrix Science, London, UK) (Perkins et al., 1999) against a combined database from a previous study (Heyer et al., 2019) consisting of UniProtKB/SwissProt and several metagenomes.

The MetaProteomeAnalyzer (version 3.1) (Heyer et al., 2019) was used for taxonomic and functional annotation of unknown protein sequences from the metagenomes by BLAST search and for grouping of redundant proteins to so-called metaproteins (protein groups) based on overlapping peptide identification. The metaproteins were mapped to their metabolic functions based on their enzyme commission and KEGG orthology numbers, using an extended pathways assignment (Heyer et al., 2019; Sikora, 2019). All mass spectrometry results were made publicly available by an upload to PRIDE (Vizcaíno et al., 2015), which could be accessed with the accession number PXD024788 (Reviewer Account: Username: Password: YKz3c6eb). More details on the protocol are provided in SI1, and the resulting tables containing the spectra (SI3), and the assigned phyla (SI4) and functions (SI5) are included in SI.

#### 2.2.3. Metabolomics

Polar to medium-polar metabolites were extracted directly from the at −80 °C frozen samples using an optimised method for *in vitro* digests (Vanden Bussche et al., 2015) with some modifications. An aliquot of 500 μL digestate was transferred to a 2 mL Eppendorf, and following the introduction of an internal standard (30 μL of a 25 ng μL^−1^ L-alanine-d3 solution), each sample was vortexed for 15 s, followed by centrifugation at 13,300*g* for 5 min at 4 °C. The supernatant was passed through a PVDF filter (0.22 μm), diluted (1:2) with ultrapure H_2_O, and transferred to a glass LC-MS (liquid chromatography – mass spectrometry) vial. A pool of extracted samples was used as a quality control standard for normalisation.

An UHPLC (ultra-high performance liquid chromatography) was achieved on a Vanquish Quaternary pumping system (Thermo Fisher Scientific, San Jose, USA), equipped with an Acquity HSS T3 column (150 × 2.1 mm, 1.8 μm) (Waters, Manchester, UK), and applying a gradient elution program. The HRMS (high resolution mass spectrometry) was performed using a Q-Exactive™ Orbitrap mass spectrometer that was operated in full-scan and polarity switching mode, across an *m/z* scan range of 53.0 – 800 Da. Methodological details are described elsewhere (De Paepe et al., 2018). Targeted data processing was carried out with Xcalibur 3.0 software (Thermo Fisher Scientific, San José, CA, USA), and used an in-house database, comprising data about > 300 metabolites (Wijnant et al., 2020). Compounds were identified based on their *m/z*-value, C-isotope profile, and retention time relative to the internal standard and authentic reference standard. Compound Discoverer™ 2.1 (Thermo Fisher Scientific, MA, USA) was used for the untargeted analysis of full-scan data, thereby applying optimized software settings (Wijnant et al., 2020). Based on the data from the negative control, noise/artefact peaks were removed. The resulting tables from untargeted (SI6) and targeted (SI7) data processing are included in the SI.

#### 2.2.4. Cytomics

Fresh samples were diluted with PBS (phosphate buffer saline), which was prepared following the instructions of the manufacturer (Sigma-Aldrich, Overijse, Belgium), to a concentration of 1 g L^−1^ TS in a 1.5 mL Eppendorf. The samples were then sonicated for 3 min at 10% amplitude on a QSonica Q700 sonicator (Qsonica L.L.C, Newtown, CT, USA) using indirect sonication with a cup horn. The samples were then directly diluted (1:1000) in 0.2 μm-filtered PBS, stained with Sybr Green I (final concentration of 1x), and incubated for 20 min at 37°C. The stained samples were analyzed on a FACSVerse flow cytometer at 60 μL min^−1^ for a maximum of 1 min, as described earlier (Props et al., 2018). The performance of the FACSVerse was verified by the FACSuite software performance quality check using CS&T research beads (BD Biosciences, San Jose, CA, USA). The raw flow cytometry data were exported in FCS format and imported into R (v3.6.1) through the *flowCore* package (v1.38.2), denoised using a gating strategy and processed with the Phenoflow package as described elsewhere (https://github.com/CMET-UGent/Phenoflow_package/wiki/1.-Phenotypic-diversity-analysis) (Props et al., 2018). The resulting table is included in the SI (SI8). The raw and denoised flow cytometry data are available from FlowRepository under ID FR-FCM-Z3RT.

### 2.3. Statistical analyses

All statistical analyses were performed in R, version 3.6.1. (http://www.rproject.org) (R Development Core Team, 2013). The α-diversity in each sample and for the four different methods was determined using the Hill diversity numbers (Hill, 1973), using the vegan package (v2.5-6). These numbers reflect the richness (H_0_), the exponential of Shannon entropy (H_1_) and the inverse Simpson (H_2_) index. The Wilcoxon signed-rank test with Bonferroni correction and Spearman and Kendall correlation tests were used to compare the α-diversity measures between the four different methods and three digester types (*i.e.*, Sludge, Agro and Dranco). The β-diversity between the different samples, based on the four methods, was determined by the non-metric multidimensional scaling (NMDS) plots. They were constructed using the Bray-Curtis distance measure (Bray and Curtis, 1957), using the phyloseq (v1.28.0), ape (v5.3) and phangorn (v2.5.5) packages. Significant differences between the digester types and relations with operational parameters were identified using pairwise permutational ANOVA (PERMANOVA) analysis (9999 permutations) with Bonferroni correction, using the *adonis* function (vegan). Multivariate homogeneity of dispersion (variance) between the different digester types was calculated using the *betadisper* function (vegan), a multivariate analogue of Levene’s test for homogeneity of variances. Significant differences were detected using the Kruskal–Wallis test and Tukey’s test for post-hoc analysis (*TukeyHSD* function). A Mantel test (*mantel* function, vegan) with Bonferroni correction, based on the Bray-Curtis dissimilarity measure and using the Spearman correlation, was used to compare the profiles of the four different methods. Canonical correspondence analysis (CCA), using the *envfit* function (vegan), was used to evaluate the strength of the correlation of the operational parameters with the bacterial community, based on the Bray-Curtis distance measure. Operational parameters with a significant impact on the microbial community profile were determined through PERMANOVA analysis (9999 permutations), and visualised through a CCA plot. Differential abundance analysis was used to identify OTUs, proteins, metabolites, or cytometric traits that showed a significant difference between the digester types using the *DESeqDataSetFromMatrix* function from the DESeq2 package (Love et al., 2014) with an adjusted *P*-value (Benjamin-Hochberg correction (Benjamini and Hochberg, 1995)) significance cut-off of 0.05 and log2 fold change cut-off of 2. The results were considered at two levels, *i.e.*, (1) the number of identified OTUs, proteins, metabolites, and cytometric traits, and (2) their total relative abundance for each method. Spearman correlation tests were used to identify correlations between the fingerprints’ features and the process parameters. For groupwise comparison between the Agro, Dranco, and Sludge digesters, Kruskal-Wallis test with Dunn posthoc test and a significance cut-off of 0.05 were applied.

### 2.4. Analytical techniques

The TS, VS and TAN were determined according to standard methods (Greenberg et al., 1992). The pH and conductivity were measured with a C532 pH and C833 conductivity meter (Consort, Turnhout, Belgium), respectively. The free ammonia (NH3) concentration was calculated based on the TAN concentration, pH and temperature in the digester. Concentrations of the Na^+^ and K^+^ cations were determined on a 761 Compact ion chromatograph (Metrohm, Herisau, Switzerland) with a Metrosep C6 e 250/4 column and Metrosep C4 Guard/4.0 guard column. The eluent contained 1.7 mM HNO3 and 1.7 mM dipicolinic acid. Sample preparation was carried out by centrifugation at 10,000*g* for 10 min, followed by filtration over a 0.45 μm filter (type PA-45/25, Macherey-Nagel, Germany) to remove all solids, and dilution with milli-Q water to reach the desired detection limits between 2 and 100 mg L^−1^, both for Na^+^ and K^+^. Concentrations of the different VFA were determined using gas chromatography, as described in SI1 (S3).

## 3. Results

### 3.1. Process parameters of the different anaerobic digesters

In this study, 64 full-scale ADs were investigated, containing continuous stirred-tank reactor (CSTR) and dry anaerobic digestion (Dranco) systems (Six and Debaere, 1992). The digesters were fed waste activated sludge, agricultural waste (manure, maize silage), food waste, and municipal solid waste (SI9). Three different digester types were considered, *i.e.*, digesters treating sludge (Sludge), digesters treating agro-industrial waste (Agro) and dry anaerobic digesters (Dranco) of which key operational parameters reflected clear differences. The Sludge digesters showed lower values for pH, temperature, total VFA, conductivity, TAN, TS and VS compared with the Agro and Dranco digesters (Table 1).

**Table 1.** Overview of the key operational parameters in the three different digester types Sludge, Agro and Dranco. Average values and standard deviations are presented. VFA = volatile fatty acids, SRT = sludge retention time, TAN = total ammonia nitrogen, COD = chemical oxygen demand.

|  | Unit | Sludge | Agro | Dranco |
| --- | --- | --- | --- | --- |
| Total VFA | mg COD L<sup>-1</sup> | 567 ± 907 | 7977 ± 9764 | 1226 ± 1258 |
| Total solids | g L<sup>-1</sup> | 43.9 ± 14.7 | 92.4 ± 27.5 | 244.5 ± 144.0 |
| Volatile solids | g L<sup>-1</sup> | 20.6 ± 8.3 | 32.6 ± 12.8 | 131.1 ± 103.5 |
| pH | - | 7.45 ± 0.09 | 7.90 ± 0.29 | 7.93 ± 0.24 |
| Temperature | °C | 34 ± 1 | 39 ± 7 | 42 ± 6 |
| SRT | days | 22 ± 7 | 48 ± 12 | 38 ± 16 |
| Conductivity | mS cm<sup>-1</sup> | 9.06 ± 1.71 | 36.74 ± 14.21 | 16.95 ± 10.85 |
| Sodium | mg L<sup>-1</sup> | 135 ± 40 | 3465 ± 2917 | 1227 ± 694 |
| Potassium | mg L<sup>-1</sup> | 245 ± 277 | 3704 ± 1715 | 1772 ± 1689 |
| TAN | mg N L<sup>-1</sup> | 1304 ± 216 | 3415 ± 1334 | 2275 ± 1314 |
| Ammonia | mg N L<sup>-1</sup> | 41 ± 17 | 391 ± 234 | 339 ± 340 |

### 3.2. Microbial community analysis

Following absolute singleton removal, the bacterial community’s amplicon sequencing resulted in an average of 46,748 ± 19156 reads per sample, representing an average of 1,142 ± 621 OTUs per sample (SI2). Metaproteomics identified on average 11,787 ± 5,995 spectra (SI3), which were assigned to 3,514 ± 1,526 metaproteins, 30 ± 4 taxonomic phyla (SI4) and 41 ± 7 core functions (SI5). Untargeted metabolome analysis revealed 16,671 components (metabolic features), with an average of 12,034 ± 2,516 components per sample (SI6). The targeted metabolomics analysis identified 47 key metabolites (SI7), with on average 43 ± 3 metabolites identified per sample. The cytomics analysis resulted in 16,920 traits, with an average of 7,264 ± 1,414 traits per sample (SI8).

#### 3.2.1. Microbial community fingerprinting: α-diversity

Microbial fingerprints were compared between the four different methods, based on α- and β-diversity. The comparison of the different α-diversity measures H_0_, H_1_, and H_2_ resulted in significant differences between the four different methods and for each of the three diversity measures (*P* < 0.0001, Wilcoxon signed-rank test, Figure 1). Exceptions were the H_0_ for the amplicon sequencing and metaproteomics (*P* = 0.74) and the H2 for the metaproteomics and metabolomics (*P* = 0.74). Spearman correlation (Table S1) analysis revealed a significant negative correlation between the amplicon sequencing and metaproteomics for H_0_ (ρ = −0.69, *P* < 0.0001), H_1_ (ρ = −0.64, *P* < 0.0001), H_2_ (ρ = −0.50, *P* < 0.0001). A significant positive correlation could be observed between the metaproteomics and metabolomics data for H_0_ (ρ = 0.47, *P* = 0.0001) and H_1_ (ρ = 0.38, *P* = 0.0026), while a negative correlation was observed between the metabolomics and cytomics for H_0_ (ρ = −0.29, *P* = 0.023), in contrast to a positive correlation for H_1_ (ρ = 0.39, *P* = 0.0021). The metaproteomics and cytomics showed a significant negative correlation for H_0_ (ρ = −0.26, *P* = 0.043), but a positive correlation for H_1_ (ρ = 0.28, *P* = 0.033) and H_2_ (ρ = 0.38, *P* = 0.0035). Similar results were obtained for the Kendall correlation analysis (Table S2). A comparison of the α-diversity in different digester types for each method revealed some clear differences (Figure 1 & Table S3). A significantly higher value for H0, H_1_ and H_2_ (*P* < 0.0001) was observed based on the amplicon sequencing data for the Sludge compared to the Agro and Dranco digesters, while there was no significant difference between the Agro and Dranco digesters for H_0_ (*P* = 1.00), H_1_ (*P* = 0.36) and H_2_ (*P* = 0.084). The opposite was true for the metaproteomics and metabolomics data, with a significantly lower H_0_ value (*P* < 0.0001) for the Sludge than the Agro and Dranco digesters. For the metaproteomics data, also H_1_ and H_2_ reached significantly higher values (*P* < 0.0001) in the Agro and Dranco digesters, compared to the Sludge digesters. The cytomics data differed from the other methods, with no significant differences between the digester types for H0. In contrast, for H1, the Dranco digesters showed significantly higher values than the Agro (*P* = 0.0006) and Sludge (*P* < 0.0001) digesters. For H2, the Dranco digesters showed the highest values again, compared to the Agro (*P* = 0.0008) and Sludge (*P* < 0.0001) digesters, but the Agro digesters also showed a significantly higher value (*P* = 0.034) than the Sludge digesters.

**Figure 1.**
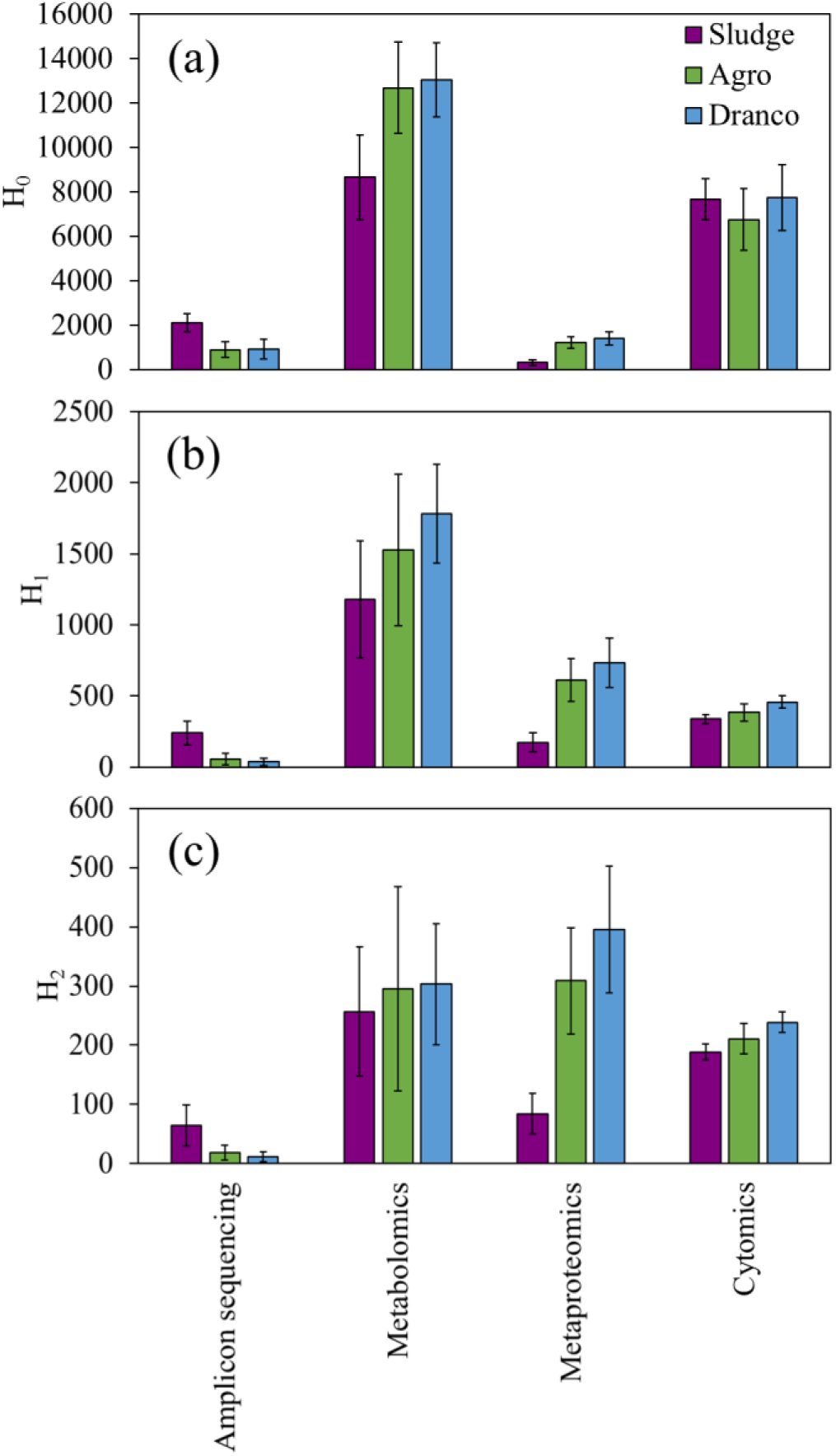
Alpha diversity, *i.e.*, the microbial community diversity within each sample, of the samples for the three digester types and four different methods, *i.e.*, amplicon sequencing (operational taxonomic units, OTUs), metabolomics (metabolites), metaproteomics (metaproteins), and cytomics (cytometric traits). The three Hill order diversity numbers (a) H_0_ (richness, number of OTUs), (b) H_1_ (exponential value of the Shannon index) and (c) H_2_ (inverse Simpson index) were calculated for each sample, and the average values are presented for each digester type for each of the four methods. Error bars represent standard deviations of average values per digester type.

#### 3.2.2. Microbial community fingerprinting: β-diversity

The β-diversity analysis, based on the Bray-Curtis dissimilarity index, revealed a significant clustering (Table S4) of the three digester types for each of the four methods, *i.e.*, based on the identified OTUs, proteins, metabolites, and cytometric traits (Figure 2). While the Sludge digesters were most clearly separated from the other digesters, reflected by the higher R^2^ values between 0.15 and 0.31 for the Sludge *vs.* Agro samples and between 0.30 and 0.42 for the Sludge *vs.* Dranco, the Agro and Dranco digesters were more closely related (R^2^ values between 0.05 and 0.13), though still significantly separated. Merging of the OTUs into their respective phyla (amplicon sequencing) and translation of the proteins into phyla or functions (metaproteomics) still resulted in a significant separation of the Sludge digesters from the other two types (Table S4, Figure S1). The separate clustering of the Agro and Dranco digesters was not possible when OTUs and proteins were combined into their respective phyla.

**Figure 2.**
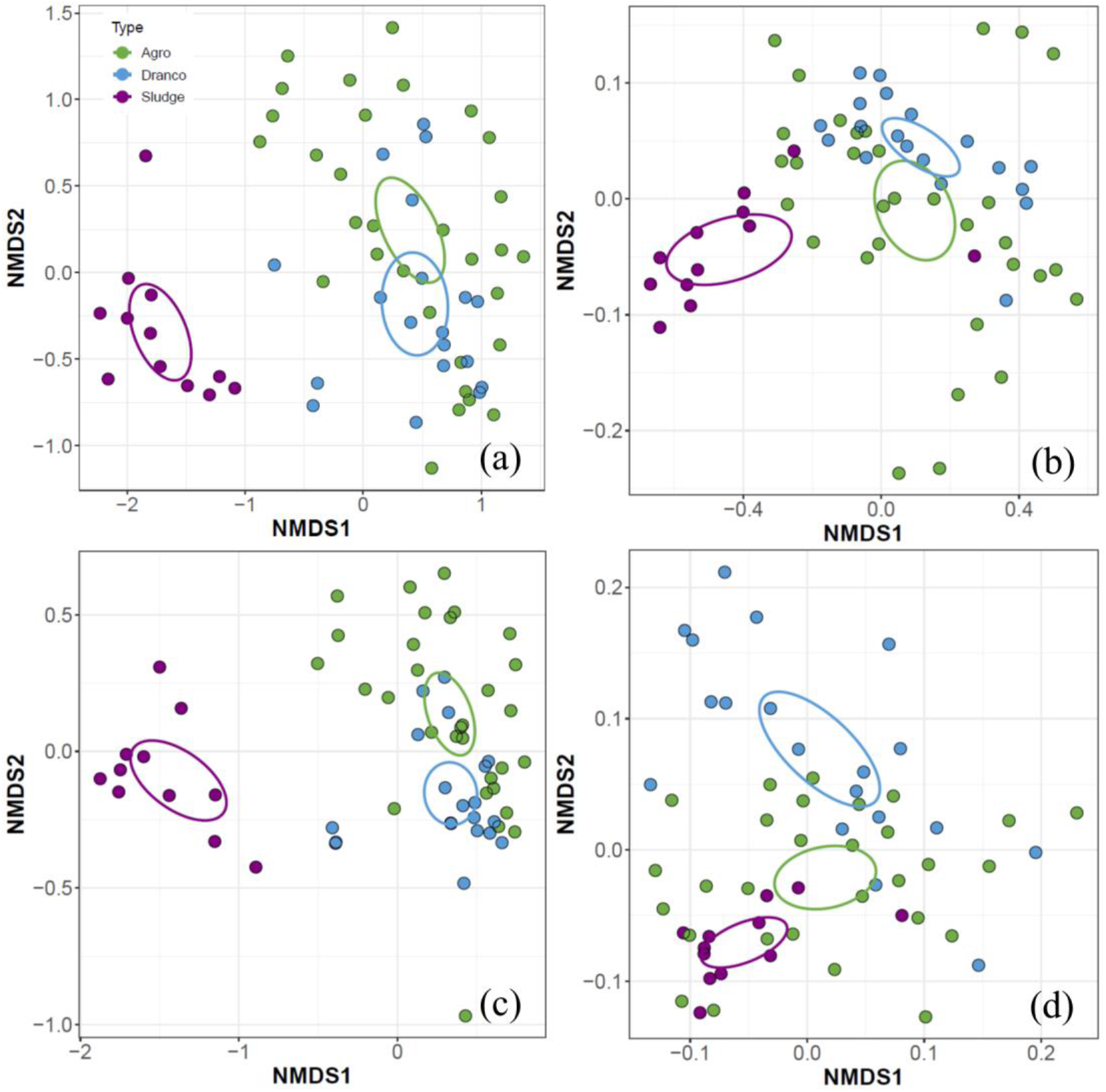
Beta diversity, *i.e.*, the difference in the microbial community between samples, based on non-metric multidimensional scaling (NMDS) analysis of the Bray-Curtis distance measure of the (a) amplicon sequencing at the OTU (operational taxonomic unit) level (stress = 0.127), (b) metabolomics (stress = 0.064), (c) metaproteomics at the metaprotein level (stress = 0.108), and (d) cytomics (stress = 0.079). The different colours represent the digester types, and the ellipses represent the 95% value of the standard error of the average value for each digester type.

The key parameters temperature, TAN, conductivity, free ammonia, total VFA and pH showed a significant relation with the amplicon sequencing, metaproteomics and metabolomics profiles (*P* < 0.05, Table S5, Figure S2), except for the free ammonia concentration and the metabolomics profile (*P* = 0.070). The cytomic profile showed a significant relation with temperature, free ammonia, total VFA and pH (*P* < 0.05, Table S5), but not with the TAN (*P* = 0.094) and conductivity (*P* = 0.072). A significantly higher degree of variance was observed in the Agro digesters compared with the Sludge digesters for the amplicon sequencing (*P* < 0.0001), metaproteomics (*P* = 0.0028), metabolomics (*P* < 0.0001) and cytomics (*P* = 0.0006) (Table S6, Figure S3). The degree of variance was also significantly higher in the Dranco digesters compared to the Sludge digesters for the amplicon sequencing (*P* = 0.0016), metabolomics (*P* = 0.0075) and cytomics (*P* = 0.0001), but not for the metaproteomics (*P* = 0.61). The degree of variance between the Agro and Dranco digesters was similar for the amplicon sequencing (*P* = 0.13) and cytomics (*P* = 0.61), but significantly different for the metaproteomics (*P* = 0.017) and metabolomics (*P* = 0.0071). The Mantel test revealed a similar β-diversity profile when comparing the amplicon sequencing with the metaproteomics (*P* = 0.0006, R^2^ = 0.86) and metabolomics (*P* = 0.0018, R^2^ = 0.24) profiles, and also when comparing the metaproteomics with the metabolomics (*P* = 0.0018, R^2^ = 0.24). In contrast, the cytomics profile did not show a significant similarity with the amplicon sequencing (*P* = 0.091, R^2^ = 0.10), metaproteomics (*P* = 0.43, R^2^ = 0.073) and metabolomics (*P* = 0.50, R^2^ = 0.078) profiles. Differential abundance analysis, which was used to identify OTUs, proteins, metabolites, or cytometric traits that show a significant difference in relative abundance between the digester types, revealed an overall stronger difference between the Sludge compared to the Agro and Dranco digesters for three of the four methods (Table 2). For the amplicon sequencing, metaproteomics and metabolomics, the percentage of OTUs, proteins and metabolites, respectively, without considering their relative abundance, that was significantly different between the Sludge vs. Agro and Dranco digesters was higher than 50% (Table 2). When their relative abundance was considered, only for the metabolomics, this value exceeded 50%, indicating that mainly low-abundant features contributed to the difference between digester types. The cytomics profile showed a similar number and total relative abundance of traits between the Agro compared to the Dranco and Sludge digesters, yet, the Sludge and Dranco digesters appeared to host less common traits, both with and without consideration of relative abundance.

**Table 2.** Differential abundance testing results. Differential abundance analysis was used to identify features (OTUs, proteins, metabolites, or cytometric traits) that showed a significant difference in relative abundance between the digester types. Average values and standard deviations are reported. An adjusted *P*-value (Benjamin-Hochberg correction) significance cutoff of 0.05 and log2 fold change cut-off of 2 were considered as thresholds. Both the percentage of changed features (OTUs, proteins, metabolites, and cytometric traits) that showed a significant difference (top) and their cumulative relative abundance (bottom) are presented. OTUs = operational taxonomic units.

| Percentage of OTUs, metabolites, proteins, and cytometric traits with a significant difference |  |  |  |  |
| --- | --- | --- | --- | --- |
|  | Amplicon sequencing | Metabolomics | Metaproteomics | Cytomics |
| Agro-Dranco | 33.7 ± 19.0 | 25.7 ± 7.6 | 24.2 ± 9.1 | 26.2 ± 1.5 |
| Agro-Sludge | 85.4 ± 10.1 | 70.7 ± 8.1 | 58.4 ± 9.6 | 22.6 ± 3.7 |
| Dranco-Sludge | 87.0 ± 8.9 | 70.4 ± 7.8 | 57.0 ± 9.0 | 39.0 ± 3.0 |

| Cumulative relative abundance of OTUs, metabolites, proteins, and cytometric traits with a significant difference |  |  |  |  |
| --- | --- | --- | --- | --- |
|  | Amplicon sequencing | Metabolomics | Metaproteomics | Cytomics |
| Agro-Dranco | 21.7 ± 3.5 | 4.4 ± 0.5 | 20.4 ± 4.0 | 23.2 ± 4.0 |
| Agro-Sludge | 41.9 ± 9.9 | 68.5 ± 3.0 | 29.5 ± 4.4 | 23.8 ± 1.9 |
| Dranco-Sludge | 41.0 ± 8.9 | 68.2 ± 3.6 | 33.8 ± 4.9 | 38.5 ± 5.4 |

#### 3.2.3. Targeted comparison of features between the different methods

The four methods showed a similar separation of the three fermenter types Sludge, Agro, and Dranco (Figure 3). However, further interpretation of the fingerprints requires an understanding of the underlying data. Here, we identified the abundant features (OTUs, metaproteins and metabolites) of the fingerprinting methods. We mreged OTUs and metaproteins to the phylum level, and metaproteins also at the functional level. Flow cytometric data were excluded for this evaluation, because they cannot be directly attributed to specific cytometric traits.

**Figure 3.**
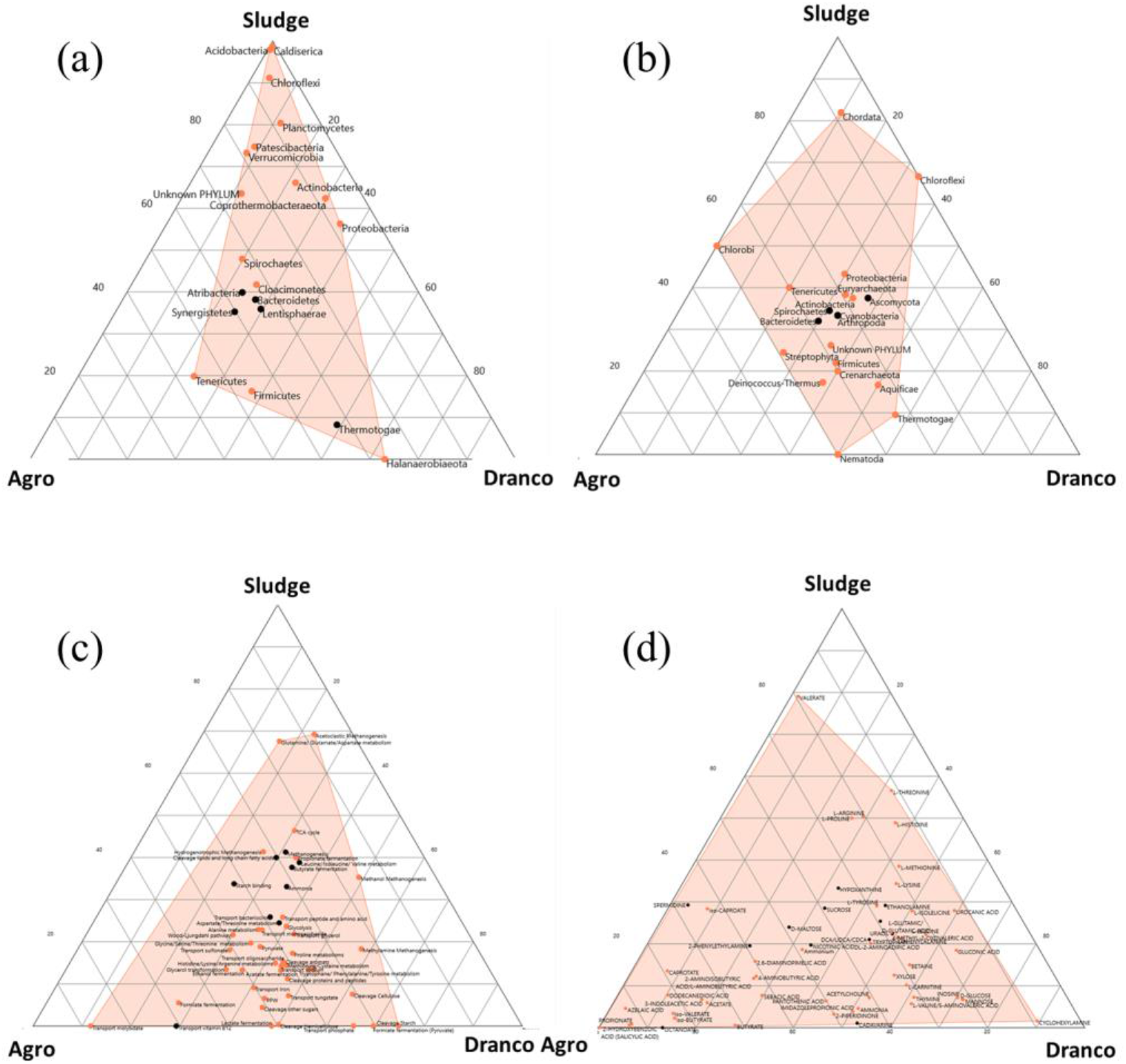
Features of the fingerprinting methods and their distribution across Sludge, Agro and Dranco digesters. The ternary plot shows the difference of the most abundant features between the Sludge, Agro and Dranco anaerobic digesters created with PAST3. It shows the ratios of the variables for the three groups as positions in an equilateral triangle. The filled region enwrap the different ratios of the variables of the three samples. The figure contains (a) the 20 top phyla identified by 16S rRNA gene amplicon sequencing, (b) the top 20 phyla identified by metaproteomics, (c) the top 48 microbial functions necessary for the anaerobic digestion process, and (d) all identified 59 metabolites (including metabolites identified via metabolomics and other operational parameters). Significant differences (Kruskal-Wallis test, *P* < 0.05) are shown as coral dots, whereas the others are shown in black.

On average, 99.2 ± 0.8% of the amplicon sequencing reads could be assigned to in total 52 bacterial phyla, with 98.5 ± 1.9% of the amplicon sequencing reads assigned to 21 phyla with an average relative abundance > 0.1% across all samples (Figure 4). The five most abundant phyla were Firmicutes (41.5 ± 19.2%), Bacteroidetes (15.8 ± 9.3%), Thermotogae (11.5 ± 16.8%), Proteobacteria (6.6 ± 7.1%), and Cloacimonetes (5.1 ± 7.6%).

**Figure 4.**
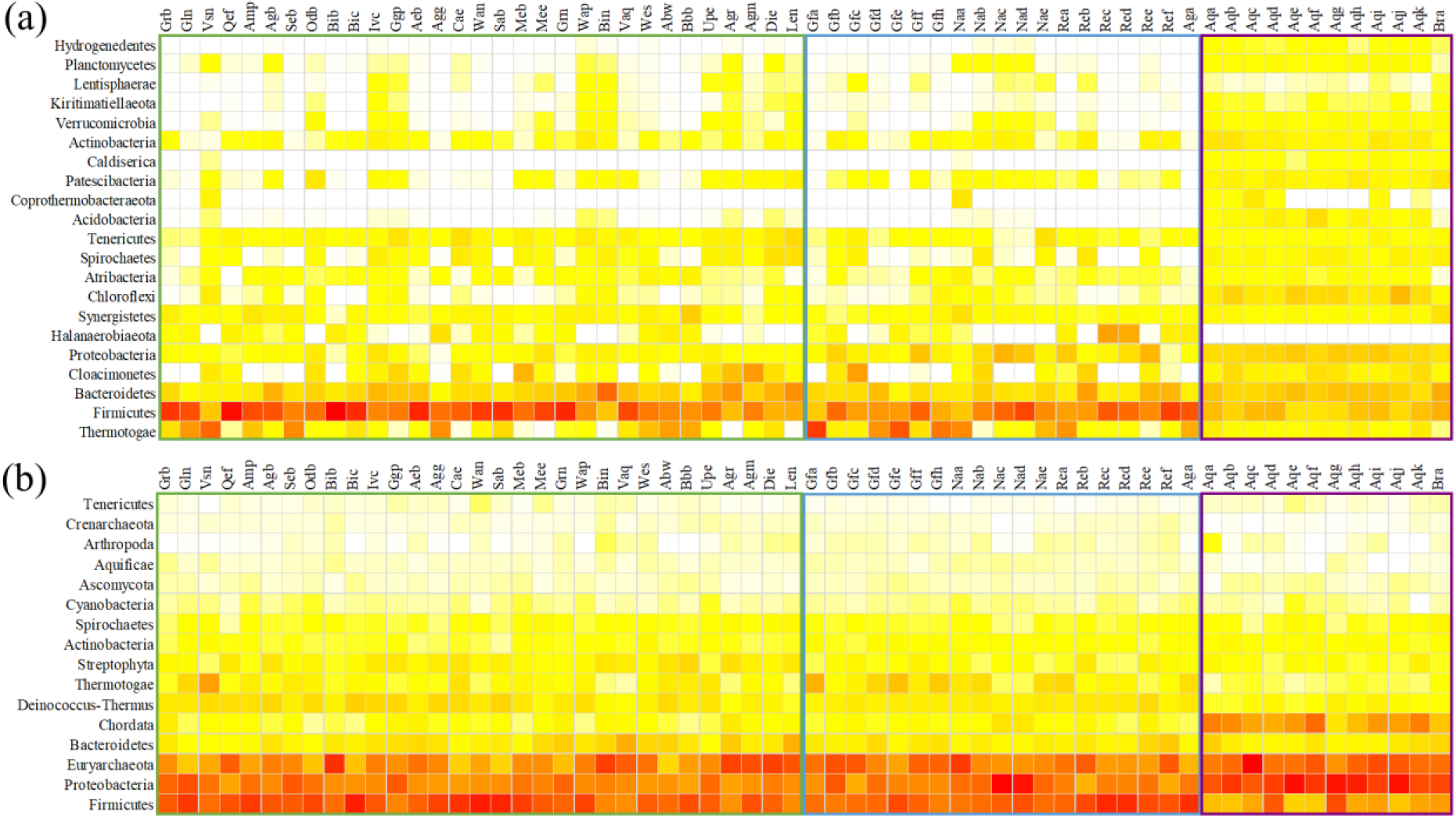
Heatmap showing the relative abundance of the microbial community at the phylum level in the Agro (left, green square), Dranco (middle, blue square) and Sludge (right, purple square) digesters based on the amplicon sequencing (a, focusing on Bacteria) and metaproteomics (b, focusing on all three Domains) data. The colour scale ranges from 0 (white) to 85% (red) relative abundance for the amplicon sequencing and 0 (white) to 30% for the metaproteomics data. Only phyla with an average relative abundance > 0.1% were included.

In contrast to the amplicon sequencing, the metaproteomics data enabled the identification of 23 bacterial, 6 archaeal and 18 eukaryotic phyla (SI4). They represented 69.1 ± 5.0% of all identified spectra (Figure 4) (based on the spectral abundance). The remaining identified spectra and metaproteins could not be assigned to any taxonomy, due to missing taxonomic annotations in the protein databases or homologous peptide identifications across the domains. When only the dominant phyla (> 0.1% of the spectral abundance) were considered, 68.5 ± 4.7% of the spectra were mapped into 9 bacterial, 2 archaeal and 5 eukaryotic phyla. The four most abundant phyla based on the spectral abundance, were Firmicutes (18.3 ± 5.1%), Proteobacteria (17.3 ± 5.0%), Euryarchaeota (15.8 ± 5.0%), Bacteroidetes (3.4 ± 2.0%), and Chordata (3.3 ± 4.6%). Eukaryotic proteins represented mainly proteins from the feedstock.

The functional metaprotein profile represented the microbial metabolism required for the anaerobic digestion process (SI5, Figure 3 & 5). It comprised 48 core functions with a cumulative abundance of 38.6 ± 3.5% of all identified spectra. On average, each digester contained 41 ± 7 functions. The most abundant functions were the final steps of methanogenesis (5.3 ± 2.5%), glycolysis (3.7 ± 1.2%), transport of oligosaccharide (3.2 ± 1.7%), hydrogenotrophic methanogenesis (2.7 ± 1.2%), and pyruvate metabolism (2.2 ± 0.9%).

**Figure 5.**
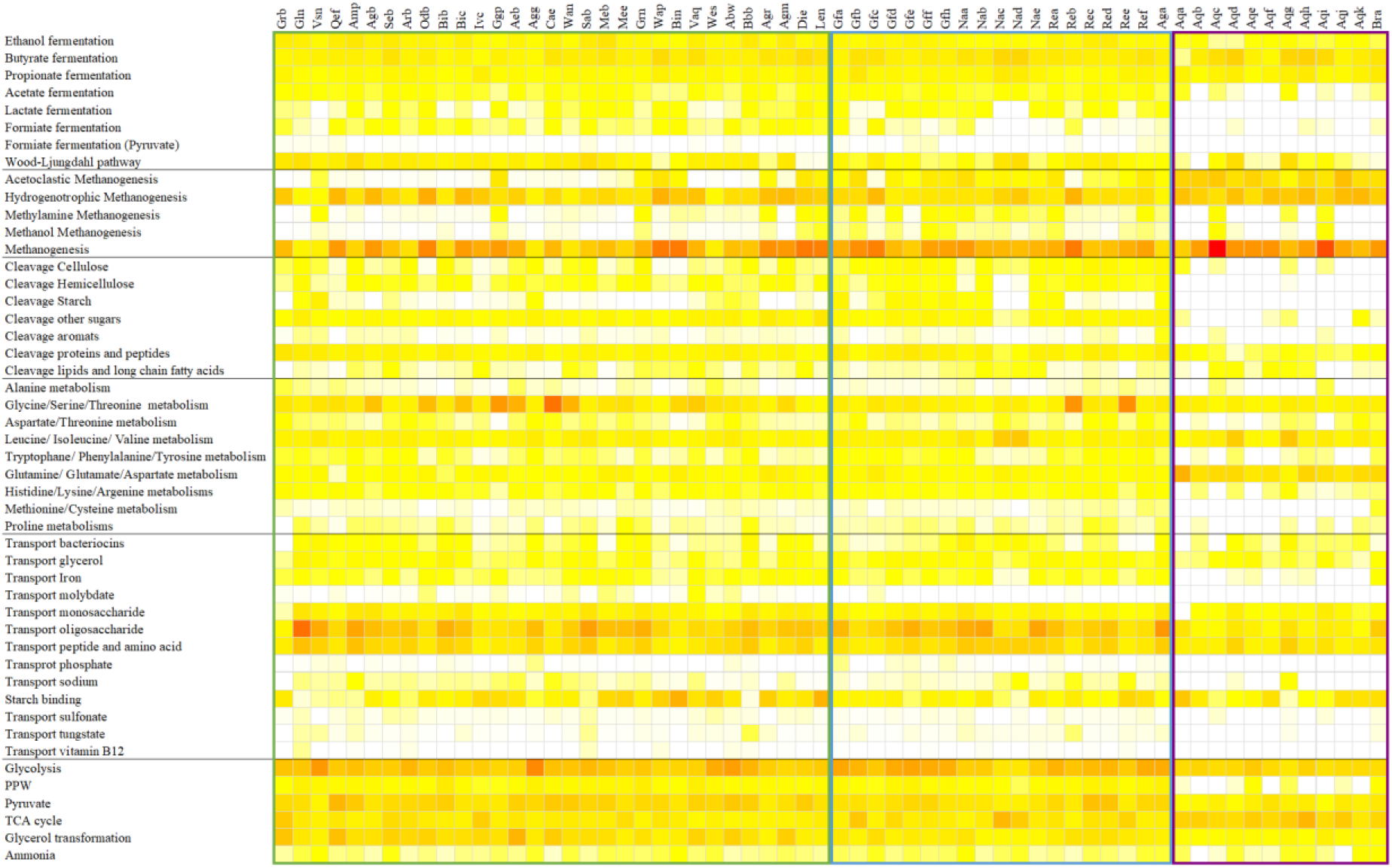
Heatmap showing the relative abundance of the different functions, based on the metaproteomics data in the Agro (left, green square), Dranco (middle, blue square) and Sludge (right, purple square) digesters. The colour scale ranges from 0 (white) to 15% (red) relative abundance. Only functions with an average relative abundance > 0.1% were included.

#### 3.2.4. Correlation within and between the features

Correlation analysis, including seven main process parameters, the three digester types, and the identified features (44 phyla for amplicon sequencing, 35 phyla for metaproteomics, 48 functions for metaproteomics, and 47 metabolites), resulted in 7,714 significant (*P* < 0.05) positive and 5,970 negative correlations (SI10). In general, similar features (*e.g.*, metaprotein phyla) possessed higher absolute correlation values among each other than to other features (SI10). Similar correlations between the phyla relative abundance to other features were observed for the amplicon sequencing and metaproteomics, *e.g.*, for Firmicutes, Bacteroidetes, Thermatogae, Chloroflexi, Actinobacteria, Proteobacteria, and Spirochaetes. Metabolic functions and their key metabolites were partially correlated, *e.g.*, glucose and glycolysis (ρ = 0.46), or acetate and acetoclastic methanogenesis (ρ = −0.61). Some correlation of functions indicated combined metabolic pathways, such as the positive correlation of the Wood-Ljungdahl pathway to ethanol fermentation (ρ = 0.56), acetate fermentation (ρ = 0.45), and glycerol transformation (ρ = 0.63). However, for most correlations, it remained unclear if they indicate a certain dependency, a preference to a certain ecological niche (*e.g.*, digester type or temperature) or false positives. Based on the *P*-value of 0.05, we would expect 684 false-positive correlations.

## 4. Discussion

Microbial community analysis of full-scale AD plants at four different levels, *i.e.*, 16S rRNA gene amplicon sequencing, metaproteomics, metabolomics, and cytomics, resulted in a different α-diversity profile. In contrast, β-diversity profiles were similar between the different methods, including their link with operational parameters, both based on the data directly or when translated into phyla or functions, which confirms the idea of triangulation. In-depth analysis of each method’s features yielded certain similar features (*e.g.*, the abundance of some phyla in the 16S rRNA gene amplicon sequencing and metaproteomics data), yet, also some clear differences.

### 4.1. Cytometric fingerprinting as a valid method to monitor the anaerobic digestion microbiome

The four different fingerprinting methods, *i.e.*, 16S rRNA gene amplicon sequencing, metaproteomics, metabolomics, and cytomics, focus on different data types and possess different resolutions, biases, and degrees of data processing. Nonetheless, each of the four methods resulted in a similar β-diversity profile, with significant separation of the three digester types, which is an excellent reflection of triangulation (Lawlor et al., 2016). Aggregation of the single features, such as OTUs or metaproteins, into higher levels, such as phyla or metabolic functions, still allowed the significant separation of the Sludge digesters from the Agro and Dranco digesters. However, the Agro and Dranco digester could no longer be separated at the phylum level, based on both the amplicon sequencing and metaproteomics data. This apparent decrease in resolution in the β-diversity profile, when considering higher phylogeny levels, can have two causes. First, numerous OTUs or metaproteins could not be assigned, even at the phylum level. Hence, they are eliminated from the dataset. Second, though relying on the most recent databases and methods, identification of taxonomy is still prone to errors (Murali et al., 2018), leading to OTUs or metaproteins wrongly assigned to a specific phylum. For example, for certain taxa, such as the recently identified Patescibacteria superphylum (Rinke et al., 2013), identified by 16S rRNA gene amplicon sequencing, only two protein sequences were available in UniProtKB/Swissprot (queried 16.02.2021). Merging of OTUs and metaproteins into taxa and functions also requires feature identification and expert knowledge, and it might suppress important features. Furthermore, the functional analysis focused on proteins relevant to anaerobic digestion but not on other functions, such as housekeeping proteins, sporulation, phage infection or adaption to environmental conditions.

In contrast to the other three methods, cytometric fingerprinting is less prone to method-inherent errors. Different steps needed to obtain a microbial community profile for amplicon sequencing, metaproteomics and metabolomics data could create several biases. Typical examples are incomplete cell lysis or an enrichment/depletion of genes/proteins/metabolites during extraction (Bonk et al., 2018; De Vrieze, 2020). In contrast, cytometric fingerprinting requires limited sample manipulation, *i.e.*, requires no cell lysis or extraction (Props et al., 2018), and almost direct measurement of the samples is possible (Frossard et al., 2016). This could explain the deviations in the cytometric fingerprint, compared to the other methods.

The amplicon sequencing, metaproteomics and metabolomics have a more direct link with process functioning. The presence of specific taxa (amplicon sequencing), proteins (metaproteomics) or metabolites (metabolomics) could explain the outcome of AD, and engineered microbial processes in more detail. For example, the presence (amplicon sequencing) and activity (metaproteomics) of certain methanogens could clarify the key pathways in the AD process (Vanwonterghem et al., 2014), and metabolomics could confirm the degradation of key metabolites (Cardona et al., 2020). Such functional information cannot be deducted from cytometric fingerprints directly. However, the high degree in similarity of the cytometric β-diversity profile compared to the other three methods in this study confirms the validity of flow cytometry as a potential microbial community fingerprinting tool in AD. Similarly, other studies confirmed the coincidence between cytometric fingerprinting and amplicon sequencing in other environments, such as freshwater ecosystems (Props et al., 2016) and landfill leachates (Kinet et al., 2016).

### 4.2. Fingerprinting or feature identification: the question determines the choice

There is an ongoing debate concerning the usage of either fingerprinting or the identification of features (taxa, proteins, metabolites) in the framework of direct microbial community-driven process monitoring in AD. Fingerprinting and the identification of features are, in fact, complementary methods (De Vrieze, 2020), and the choice for either the one or the other approach will depend on whether the goal is overall community or specific function monitoring. There are, however, several challenges that hamper the application of microbial community monitoring in AD or other engineered microbial processes.

The first challenge is the analysis speed. Flow cytometry has the advantage that it can be applied for online fingerprinting, as demonstrated in drinking water production (Buysschaert et al., 2018; Favere et al., 2020) and axenic culture bioproduction processes (Broger et al., 2011). Thus, a cytometric fingerprint can be obtained in less than an hour, yet, this remains, at present, elusive in more complex systems, such as AD. In contrast, amplicon sequencing or other “omics” methods require a comprehensive sample preparation that takes at least one day of analysis time (Heyer et al., 2019).

The second challenge is information gain. The application of fingerprinting techniques will only be beneficial if they add novel information to the operators compared to conventional process parameters, *e.g.*, methane production, pH, TAN, or VFA. The desired knowledge could include information about microbial interactions (Heyer et al., 2019) or the activity of the methanogenic Archaea or hydrolytic Bacteria (Munk et al., 2012; Rettenmaier et al., 2020).

The third challenge reflects the consistency between methods, and here fingerprinting has an advantage over identifying features. Fingerprinting in AD appears to be consistent across the different methods when one considers β-diversity, even when data are merged at higher levels of phylogeny, although to a lesser extent. This observation matches other studies comparing amplicon sequencing with T-RFLP (De Vrieze et al., 2018; Goux et al., 2015). However, when α-diversity, especially richness, is considered, there is a significant inconsistency between the different methods, which inherently relates to the depth, *i.e.*, the (relative) abundance of features that are still observed, and applied normalisation methods (De Vrieze, 2020).

### 4.3. Extending our knowledge about microbiomes and fingerprints

As shown in previous studies (De Vrieze et al., 2016; Heyer et al., 2013; Kirkegaard et al., 2017; Repinc et al., 2018), microbial fingerprints enable monitoring of the AD process’ stability, and also single features of the fingerprinting methods correlate with process parameters. For example, the phylum Chloroflexi prefers the conditions in anaerobic digesters treating sewage sludge as feedstock (Bovio-Winkler et al., 2021; Speirs et al., 2019). Several correlations could be identified between TAN, temperature, feedstock composition, and digester types with taxa and metaproteins identified in previous studies (De Vrieze et al., 2015; Heyer et al., 2016; Zhang et al., 2014). However, prediction of future process performance or shifts in the operation based on fingerprints, remains, at present, impossible. Reasons are the microbiome complexity, the lack of knowledge about the microbiome, and the phenomenon that for similar process conditions different microbiome compositions exist that may achieve a comparable process performance (Kohrs et al., 2017).

An example of the complexity and our lack of knowledge is reflected in syntrophic interactions. A strong correlation was observed between bacterial acetate fermentation, ethanol fermentation, and glycerol metabolism, matching the described pathways for syntrophic ethanol degradation in a co-culture experiment (Keller et al., 2019). Glycerol metabolism correlated to this pathway, since it comprised the missing alcohol dehydrogenase to combine both pathways. In this study, the identified NADP-dependent isopropanol dehydrogenase (Meta-Protein 2708) had even the highest spectral count of all identified proteins (SI3), emphasizing the potential importance of this new pathway. On the downside, the imprecise assignment to glycerol metabolism and a proper taxonomy (in our study, the taxonomy was *Entamoeba histolytica*) shows the requirement for better databases and further isolation and characterization of single species.

Another example of the difficulty of feature interpretations is the issue that features can have multiple functions. Enzymes for glutamine, glutamate and aspartate metabolism were abundantly present in Sludge digesters, and correlated with low values for pH and TAN. The reason could be (1) elevated amino acid metabolism or (2) nitrogen assimilation (Bernard and Habash, 2009). Glutamate is also required for osmotic regulation, since it is a counterion for potassium, only present in Sludge digesters at low concentrations (Yan, 2007) (Table 1 & SI3). A strategy to utilize the omics data for improved process operation of AD would be to shift from fingerprinting methods to panel development. Analogous to clinical panels, such as the blood panel, the panels would summarize the omics methods’ features (*e.g.*, the abundance of methanogens). Subsequently, we have to define safe operation borders for these panels by extensive experiments and modelling, before providing them to AD operators. For example, biogas production and process stability can be monitored based on the abundance of methanogens and their enzymes (Heyer et al., 2013; Munk et al., 2012), and a multivariate integrative method, based on a microbial signature of AD inhibition by ammonia and phenol could be used to predict ammonia inhibition (Poirier et al., 2020). On-demand, the complexity of these panels could be extended. Besides the abundance of methanogens and their enzymes, it might be beneficial to add thermodynamic considerations (*i.e.*, process temperature and metabolite concentrations) or the abundance of trace elements, co-factors, and hydrolytic enzymes.

## 5. Conclusions

The four different fingerprinting methods, *i.e.*, based on amplicon sequencing, metaproteomics, metabolomics and cytomics, revealed a similar clustering of the microbiomes in AD. This finding highlights that cytometric fingerprinting, a novel approach in AD, concerns a valid method for fast microbiome fingerprinting in AD, which is a key advantage over other methods. The information provided through fingerprinting can have its merits for process stability monitoring, and the identification of key features, either taxa, proteins or metabolites, gives rise to a more direct interpretation of process performance. To provide this knowledge to AD operators, new strategies are required. We propose that the microbiomes’ key features should be summarized into panels and linked with guidance for the operators.

## Acknowledgments

Jo De Vrieze was supported as postdoctoral fellow by the Research Foundation Flanders (FWO-Vlaanderen). Robert Heyer is supported by the German Federal Ministry of Education and Research (de.NBI network, project MetaProtServ, grant no. 031L0103). Ruben Props was supported as postdoctoral fellow by the Research Foundation Flanders (FWO) under grant 1221020N. Nico Boon was supported by the Bijzonder Onderzoeksfonds of Ghent University (BOF15/GOA/006). The authors would like to thank Tim Lacoere and Corina Siewert for their laboratory support. Innolab, Aquafin and OWS are kindly acknowledged for contributing to the sample collection.

## Notes

### Competing Interest Statement

The authors have declared no competing interest.

