## Supplementary file 1 for "Triangulation of microbial fingerprinting in anaerobic digestion reveals consistent fingerprinting profiles"

### S1. Amplicon sequencing and data processing

S1.1. Amplicon sequencing

Amplicon sequencing was carried out on the Illumina Miseq platform using V3 chemistry. The primers 341F (5'- NNNNNNNNNTCCTACGGGNGGCWGCAG) and 785R (5'- NNNNNNNNNNTGACTACHVGGGTATCTAAKCC) that target the V3-V4 region of the 16S rRNA gene (Klindworth et al., 2013), with an extra wobble position in the reverse primer to make it more universal, were used to target total bacteria. The DNA extracts had a concentration of 1 ng $\mu L^{-1}$, and were free of RNA, as validated with agarose gel electrophoresis. The PCR mix included 5 ng of DNA, 15 pmol of both the forward primer 341F and reverse primer 785R in 20 µL volume of MyTaq buffer containing 1.5 units MyTaq DNA polymerase (Bioline) and 2 µL of BioStabII PCR Enhancer (Sigma). For each sample, the forward and reverse primers had the same unique 10-nt barcode sequence. The PCR run contained 20-25 cycles, using the following parameters: 2 min 96 °C pre-denaturation; 96 °C for 15 s, 50 °C for 30 s, 70 °C for 90 s. Next, about 20 ng amplicon DNA of each sample were pooled for up to 48 samples carrying different barcodes (Nextera XT index kit, Illumina). The amplicon pools were purified using Ampure XP beads to remove primer dimers and other small mispriming fragments, according to the manufacturer's instructions, and their size checked on a Fragment analyser (Advanced Analytical Technologies, Inc., Ames, Iowa, USA), and quantified by fluorometric analysis. Next, multiplexing, clustering and sequencing was carried out on an Illumina MiSeq with the paired-end (2×) 300 bp protocol and indexing. The sequencing run was analysed *via* the Illumina CASAVA pipeline (v1.8.3) in which demultiplexing was based on sample-specific barcodes. The raw sequencing data were processed, removing sequence reads of too low quality (only "passing filter" reads were selected), and discarding reads containing adaptor sequences or PhiX control with an in-house filtering protocol. A quality assessment on the remaining reads was performed using the FASTQC quality control tool version 0.10.0.

S1.2. Data processing

The Mothur software package (v.1.42.3), and guidelines (Schloss et al., 2009) were used to process the raw Illumina data on a GNU/Linux 3.16.0-46-generic x86_64 system. The forward and reverse reads were assembled into contigs by a heuristic approach, taking the Phred quality scores into account. Ambiguous contigs or with unsatisfying overlap were removed, and the remaining sequences were aligned to the mothur formatted silva seed v132 database. Those sequences not aligning within the region targeted by the primer set or sequences with homopolymer stretches with a length > 12 bp were removed. The sequences were pre-clustered, allowing mismatch for every 100 bp of sequence. Chimeric sequences were removed with UCHIME (Edgar et al., 2011). Classification of the sequences was carried out by a naïve Bayesian classifier, using the RDP 16S rRNA gene training set, release 16, with an 85% cut-off for the pseudobootstrap confidence score. Taxa that were annotated as Chloroplast, Mitochondria, unknown, Archaea or Eukarya at the kingdom level were excluded. Sequences were clustered into OTUs (operational taxonomic units) with an average linkage, and at a 97% sequence identity, using the OptiClust method (Westcott and Schloss, 2017). Representative sequences were picked for each OTU as the most abundant sequence within that OTU.

### S2. Metaproteomics

Metaproteome analysis comprised phenol extraction in a ball mill, amido black protein assay, FASP digestion and LC-MS/MS measurement using a timsTOF™ mass spectrometer (Bruker

Daltonik GmbH, Bremen). A detailed description of the laboratory workflow could be found in (Robert Heyer et al., 2019). For the LC-MS/MS measurement 5 µL of each sample (1.5 µg) were injected and separated by UltiMate® 3000 nano splitless reversed-phase nanoHPLC (Thermo Fisher Scientific, Dreieich) equipped with a reversed-phase trap column (nano trap cartridge, 300 µm i.d. x 5 mm, packed with Acclaim PepMap100 C18, 5 µm, 100 Å, nanoViper, Bremen, Germany) and a reversed-phase separation column (Acclaim PepMap RSLC, C18, 2 µm, 75 µm, 50 cm, Bremen, Germany). The gradient was 5 % to 35 % mobile phase B (acetonitrile, 0.1 % formic acid) over 120 min at a flow rate of 0.4 µL $min^{-1}$. The LC was coupled directly to the MS. The timsTOF™ mass spectrometer (Bruker Daltonik GmbH, Bremen) was equipped with a captive spray ionization (CSI) source operated in positive ion mode with a capillary voltage of 1600 V. PASEF scan mode was performed using trapped ion mobility spectrometry (TIMS) from a range of 0.6 to 1.6 V*s/cm2, charge state from 0 to 5 and 10 PASEF MS/MS scans per cycle. MS/MS data acquisition was performed over the mass range from 100 to 1,700 m/z using collision-induced dissociation (CID) with an active exclusion of the same precursor after 1 spectrum for 24 seconds or a release if the ratio between intensity/previous intensity exceeded 4.

Compass DataAnalysis 5.1 software (Bruker Daltonik GmbH, Bremen) was used to process the results files of the timsTOF™ MS/MS measurement and to create mascot generic files (.mgf) for the subsequent protein identification using Mascot™ 2.6.1 (Matrix Science, London) (Perkins et al., 1999). All protein database searches used the following parameters: enzyme trypsin, one missed cleavage, monoisotopic mass, carbamidomethylation (cysteine) as a fixed modification, oxidation (methionine) as variable modifications, ± 0.02 Da precursor and ± 0.02 Da MS/MS fragment tolerance, 1 $^{13}C$ and +2/+3 charged peptide ions. As a protein database, a combined database from a previous study (R. Heyer et al., 2019) was used consisting of UniProtKB/SwissProt and several metagenomes. The result files of the mascot search were uploaded into the MPA software version 3.1 (Robert Heyer et al., 2019). A false discovery rate of 1 % was applied for the searches and identified proteins without taxonomic and functional classification were annotated with UniProtKB metadata by using protein BLAST (NCBI-Blast-version 2.2.31 (Altschul et al., 1990; Camacho et al., 2009) against the UniProtKB/SwissProt database using an e-value cutoff of $10^{-4}$. Afterward, all protein BLAST proposals with the best e-value were merged and used to annotate a protein.

Redundant, homologous protein identifications were grouped to so-called metaproteins (protein groups), when they share at least one overlapping peptide identification. Finally, a matrix of all metaproteins over all samples was exported as comma separated file containing the spectral count for each sample and also the following annotation information: NCBI taxonomy, enzyme commission numbers (EC), KEGG orthologies (KO), the UniProtKB reference clusters and the UniProtKB keywords. All MS results were made available to the public by an upload to PRIDE (Vizcaíno et al., 2015), which could be accessed with the accession number PXD024788 (Reviewer Account: Username: Password: YKz3c6eb).

#### S3. Volatile fatty acids (VFA) analysis

The VFA concentrations (C2-C8) were measured with a gas chromatograph (GC-2014, Shimadzu®, The Netherlands), equipped with a DB-FFAP 123-3232 column (30 m × 0.32 mm × 0.25 µm; Agilent, Belgium) and a flame ionization detector (FID). The samples (2 mL) were conditioned with sulphuric acid and sodium chloride, and 2-methyl hexanoic acid was used as internal standard to quantify the extraction with diethyl ether. The extracted sample (1 µL) was injected at 200 ºC with a split ratio of 60 and a purge flow of 3 mL $min^{-1}$. The oven temperature increased with 6 ºC $min^{-1}$ from 110 ºC to 165 ºC, where it was kept for 2 min. The FID had a temperature of 220 ºC. The carrier gas was $N_2$ at a flow rate of 2.49 mL $min^{-1}$. The detection limit was 30 mg $L^{-1}$ for acetate, 10 mg $L^{-1}$ for propionate and 2 mg $L^{-1}$ for the other VFA (C4-C8).

#### S4. Microbial community fingerprinting

**Table S1** Spearman correlation values for the Hill diversity numbers (Hill, 1973). These numbers reflect the richness ($H_0$), the exponential of Shannon entropy ($H_1$) and the inverse Simpson ($H_2$) index. The ρ values are presented with the P-values between brackets.

| | | Amplicon sequencing | Metabolomics | Metaproteomics |
|---|---|---|---|---|
| $H_0$ | Metabolomics | -0.46 (0.0002) | | |
| | Metaproteomics | -0.69 (<0.0001) | 0.47 (0.0001) | |
| | Cytomics | 0.49 (<0.0001) | -0.29 (0.023) | -0.26 (0.043) |
| $H_1$ | Metabolomics | -0.23 (0.077) | | |
| | Metaproteomics | -0.64 (<0.0001) | 0.38 (0.0026) | |
| | Cytomics | -0.15 (0.25) | 0.39 (0.0021) | 0.28 (0.033) |
| $H_2$ | Metabolomics | -0.020 (0.88) | | |
| | Metaproteomics | -0.50 (<0.0001) | 0.17 (0.18) | |
| | Cytomics | -0.24 (0.068) | 0.20 (0.12) | 0.38 (0.0035) |

**Table S2** Kendall correlation values for the Hill diversity numbers (Hill, 1973). These numbers reflect the richness ($H_0$), the exponential of Shannon entropy ($H_1$) and the inverse Simpson ($H_2$) index. The τ values are presented with the P-values between brackets.

| | | Amplicon sequencing | Metabolomics | Metaproteomics |
|---|---|---|---|---|
| $H_0$ | Metabolomics | -0.32 (0.0002) | | |
| | Metaproteomics | -0.50 (<0.0001) | 0.33 (0.0001) | |
| | Cytomics | 0.33 (0.0002) | -0.20 (0.022) | -0.17 (0.061) |
| $H_1$ | Metabolomics | -0.14 (0.12) | | |
| | Metaproteomics | -0.45 (<0.0001) | 0.25 (0.0042) | |
| | Cytomics | -0.10 (0.26) | 0.28 (0.0018) | 0.19 (0.038) |
| $H_2$ | Metabolomics | -0.011 (0.90) | | |
| | Metaproteomics | -0.32 (0.0002) | 0.10 (0.24) | |
| | Cytomics | -0.16 (0.068) | 0.14 (0.13) | 0.26 (0.0041) |

**Table S3** Statistical comparison of the Hill diversity numbers (Hill, 1973) between the different digester types using the Wilcoxon signed-rank test with Bonferroni correction. These numbers reflect the richness ($H_0$), the exponential of Shannon entropy ($H_1$) and the inverse Simpson ($H_2$) index. The P-values following Bonferroni correction are presented for each comparison.

| | | $H_0$ | $H_1$ | $H_2$ |
|---|---|---|---|---|
| Amplicon sequencing | Agro-Dranco | 1 | 0.36 | 0.084 |
| | Agro-Sludge | <0.0001 | <0.0001 | <0.0001 |
| | Dranco-Sludge | <0.0001 | <0.0001 | <0.0001 |
| Metabolomics | Agro-Dranco | 1 | 0.41 | 1 |
| | Agro-Sludge | <0.0001 | 0.23 | 1 |
| | Dranco-Sludge | 0.0001 | 0.0012 | 0.49 |
| Metaproteomics | Agro-Dranco | 0.045 | 0.021 | 0.0093 |
| | Agro-Sludge | <0.0001 | <0.0001 | <0.0001 |
| | Dranco-Sludge | <0.0001 | <0.0001 | <0.0001 |
| Cytomics | Agro-Dranco | 0.058 | 0.0006 | 0.0008 |
| | Agro-Sludge | 0.092 | 0.085 | 0.034 |
| | Dranco-Sludge | 1 | <0.0001 | <0.0001 |

**Table S4** Statistical comparison of the bacterial community, based on the Bray-Curtis (Bray and Curtis, 1957) distance measure between in the different digester types through pairwise permutational ANOVA (PERMANOVA) analysis (9999 permutations) with Bonferroni correction, using the *adonis* function (vegan). The P-values following Bonferroni correction are presented for each comparison.

| | Agro-Dranco | Agro-Sludge | Dranco-Sludge |
|---|---|---|---|
| Amplicon sequencing (OTUs) | 0.011 | 0.0003 | 0.0003 |
| Amplicon sequencing (Phyla) | 0.1005 | 0.0003 | 0.0003 |
| Metabolomics | 0.010 | 0.0003 | 0.0003 |
| Metaproteomics (proteins) | 0.0015 | 0.0003 | 0.0003 |
| Metaproteomics (phyla) | 0.13 | 0.0003 | 0.0003 |
| Metaproteomics (function) | 0.048 | 0.0003 | 0.0003 |
| Cytomics | 0.0033 | 0.0063 | 0.0003 |

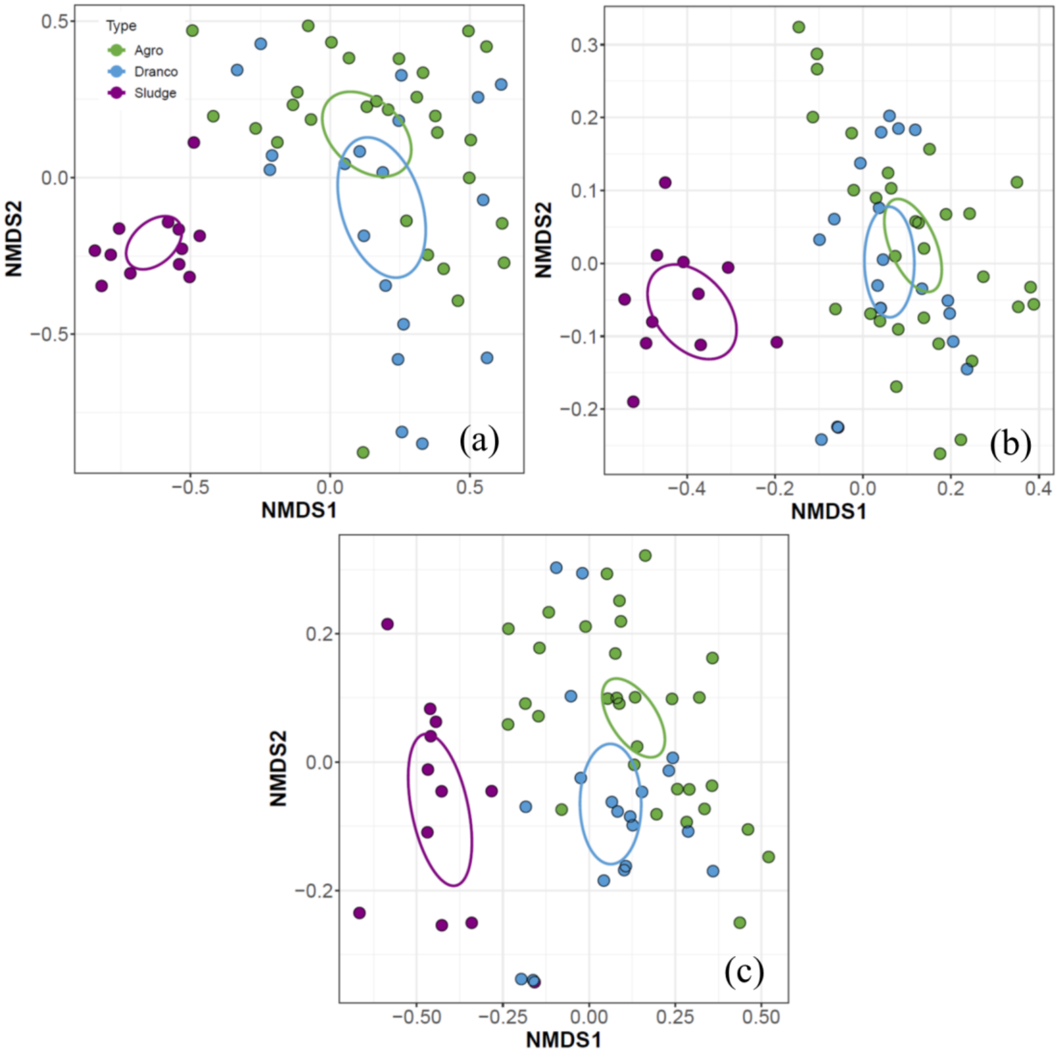

**Figure S1** Non-metric multidimensional scaling (NMDS) analysis of the Bray-Curtis distance measure of the (a) amplicon sequencing at the phylum level (stress = 0.138), (b) metaproteomics at the phylum level (stress = 0.116), and (c) metaproteomics at the function level (stress = 0.137). The different colours represent the digester types, and the ellipses represent the 95% value of the standard error of the average value for each digester type.

**Table S5** Statistical evaluation of the relation between different operational parameters and the bacterial community, based on the Bray-Curtis (Bray and Curtis, 1957) permutational ANOVA (PERMANOVA) analysis (9999 permutations), using the *adonis* function (vegan). The *P*-values are presented for all parameters for each of the four methods. TAN = total ammonia nitrogen, VFA = volatile fatty acids

| | Amplicon sequencing | Metabolomics | Metaproteomics | Cytomics |
|---|---|---|---|---|
| Temperature | 0.0006 | 0.0024 | 0.001 | 0.0016 |
| TAN | 0.0001 | 0.0001 | 0.0002 | 0.094 |
| Conductivity | 0.017 | 0.0056 | 0.0054 | 0.072 |
| Free ammonia | 0.028 | 0.070 | 0.035 | 0.0056 |
| VFA | 0.0007 | 0.0004 | 0.0017 | 0.036 |
| pH | 0.0001 | 0.0001 | 0.0001 | 0.029 |

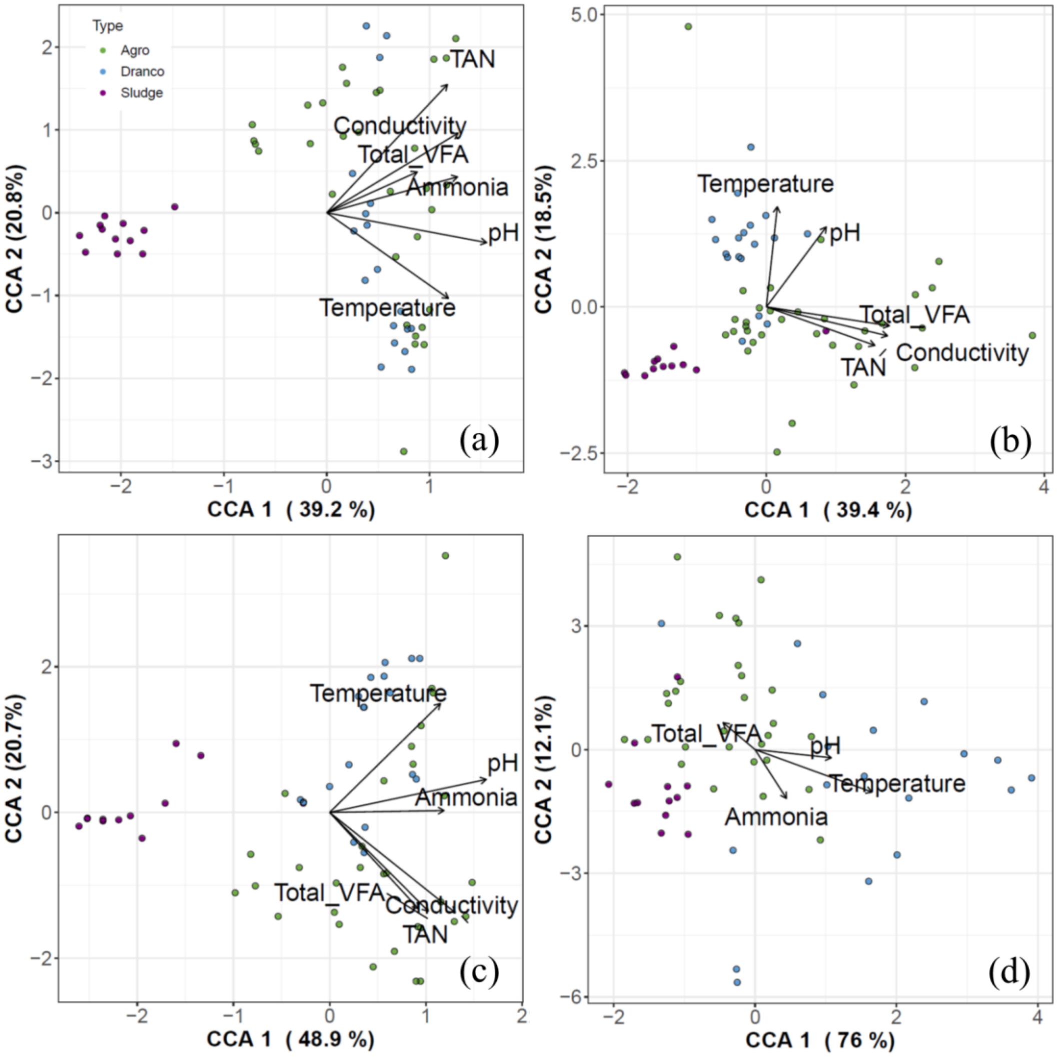

**Figure S2** Canonical correspondence analysis (CCA) of the (a) amplicon sequencing at the OTU (operational taxonomic unit) level, (b) metabolomics, (c) metaproteomics, and (d) cytomics profile. The PERMANOVA analysis (9999 permutations) identified the relationship between each profile and the operational parameters, and significant ($P < 0.05$, Table S5) correlations are presented by the arrows. TAN = total ammonia nitrogen, Total VFA = total volatile fatty acids. Ammonia reflects the free ammonia ($NH_3$) concentration.

**Table S6** Statistical evaluation of the homogeneity of variances between the different digester types for the four different methods, based on the Bray-Curtis (Bray and Curtis, 1957) distance measure. The *betadisper* function (vegan) was used to determine the distance from the centroid for all digesters in each of the three types after which the Kruskal–Wallis test with Tukey's test for post-hoc analysis (*TukeyHSD* function) was used for comparison of the different digester types. The P-values of the Tukey's test are presented for each comparison.

| | Amplicon sequencing | Metabolomics | Metaproteomics | Cytomics |
|---|---|---|---|---|
| Agro-Dranco | 0.13 | 0.0071 | 0.017 | 0.61 |
| Agro-Sludge | <0.0001 | <0.0001 | 0.0028 | 0.0006 |
| Dranco-Sludge | 0.0016 | 0.0075 | 0.61 | 0.0001 |

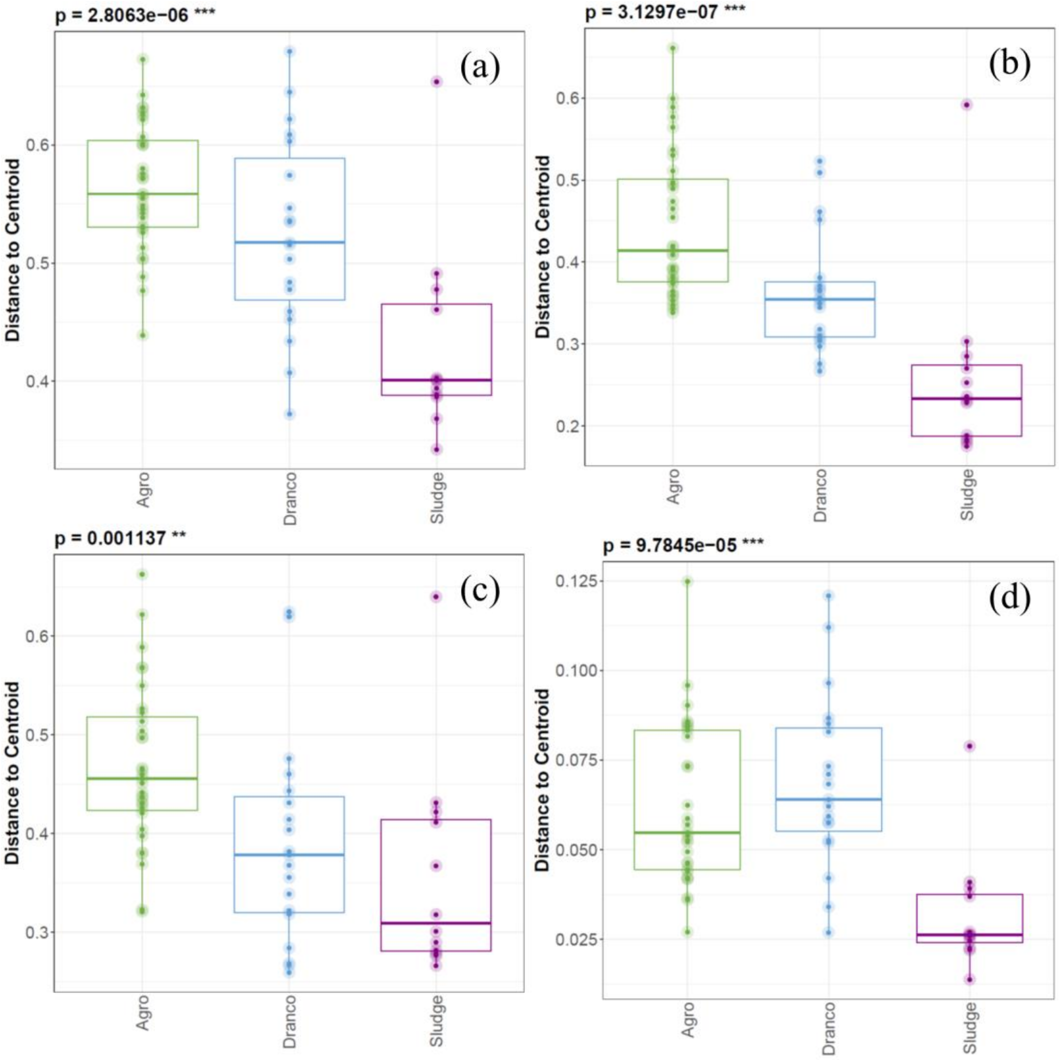

**Figure S3** Variance between the different digester types, determined as the distance from the centroid (average value) of the principal coordinates of the Bray-Curtis (Bray and Curtis, 1957) distance measure, for the (a) amplicon sequencing at the OTU (operational taxonomic unit) level, (b) metabolomics, (c) metaproteomics, and (d) cytomics profile. The P-values represent the outcome of the Kruskal–Wallis test, and the results of the Tukey's tests for post-hoc analysis are presented in Table S6.

#### S5. Bacterial community composition

Here, an overview of the bacterial community composition at the different phylogenetic levels, *i.e.*, phylum, class, order, family and OTU, is presented for all samples in the three digester types.

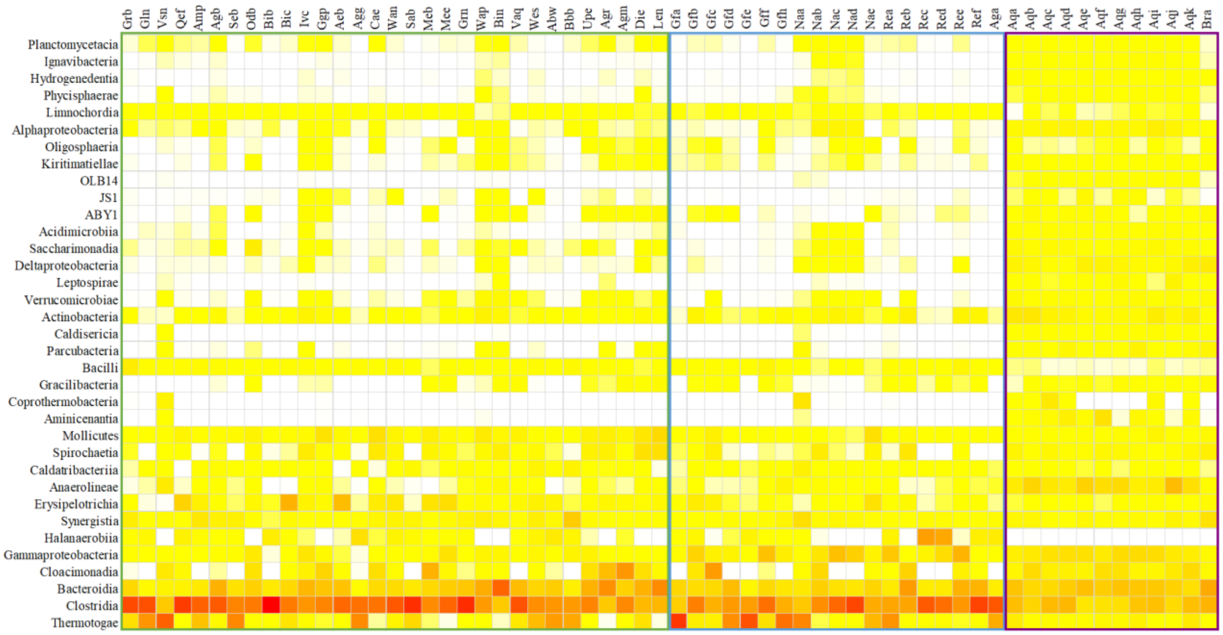

**Figure S4** Heatmap showing the relative abundance of the bacterial community at the class level in the Agro (left, green square), Dranco (middle, blue square) and Sludge (right, purple square) digesters. The colour scale ranges from 0 (white) to 90% (red) relative abundance.

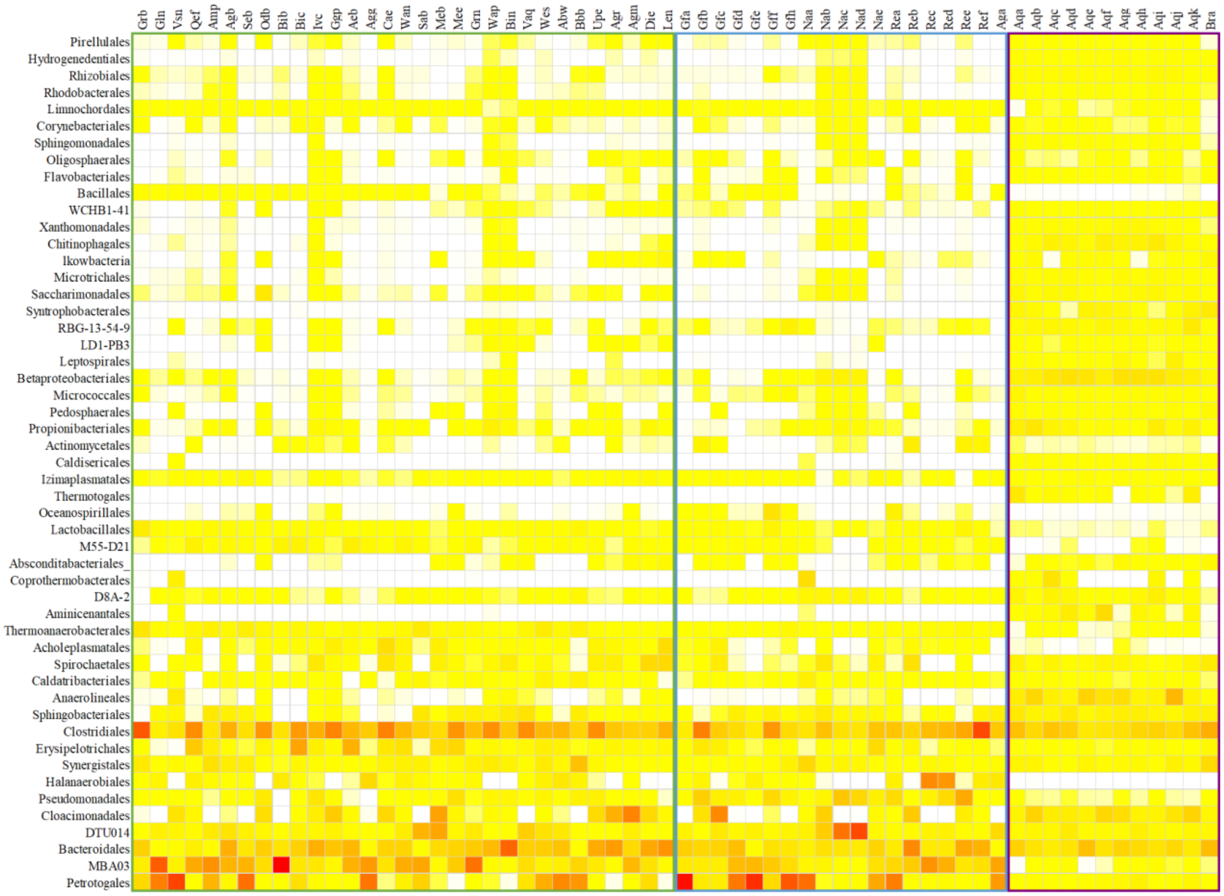

**Figure S5** Heatmap showing the relative abundance of the bacterial community at the order level in the Agro (left, green square), Dranco (middle, blue square) and Sludge (right, purple square) digesters. The colour scale ranges from 0 (white) to 70% (red) relative abundance.

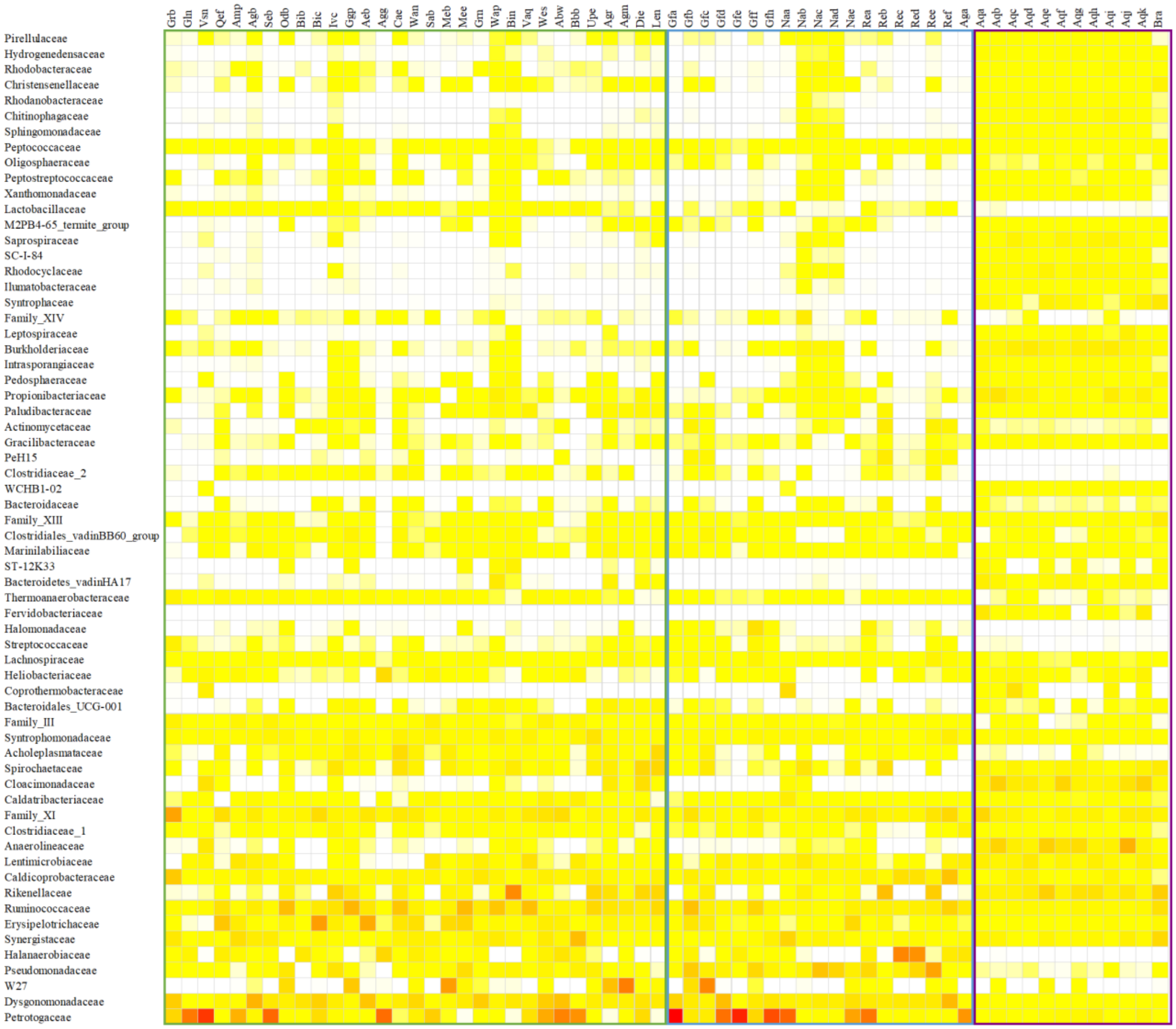

**Figure S6** Heatmap showing the relative abundance of the bacterial community at the family level in the Agro (left, green square), Dranco (middle, blue square) and Sludge (right, purple square) digesters. The colour scale ranges from 0 (white) to 70% (red) relative abundance.

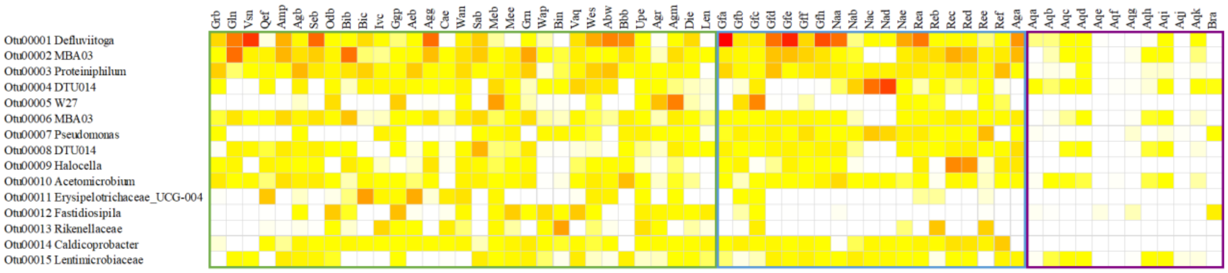

**Figure S7** Heatmap showing the relative abundance of the bacterial community at the OTU (operational taxonomic unit) level in the Agro (left, green square), Dranco (middle, blue square) and Sludge (right, purple square) digesters. The colour scale ranges from 0 (white) to 70% (red) relative abundance.
